# Inter-chromosomal linkage disequilibrium and linked fitness cost loci associated with selection for herbicide resistance

**DOI:** 10.1101/2021.04.04.438381

**Authors:** Sonal Gupta, Alex Harkess, Anah Soble, Megan Van Etten, James Leebens-Mack, Regina S Baucom

**Affiliations:** Ecology and Evolutionary Biology Department, 4034 Biological Sciences Building, University of Michigan, Ann Arbor, MI 48109; Department of Crop, Soil, and Environmental Sciences, Auburn University, Auburn, AL 36849; HudsonAlpha Institute for Biotechnology, Huntsville, AL 35806; Biology Department, Pennsylvania State University, Dunmore, PA 18512; Department of Plant Biology, University of Georgia, Athens, GA, 30602

**Keywords:** NTSR resistance, detoxification, cost, interchromosomal linkage disequilibrium, genetic hitchhiking

## Abstract

The adaptation of weedy plants to herbicide is both a significant problem in agriculture and a model for the study of rapid adaptation under regimes of strong selection. Despite recent advances in our understanding of simple genetic changes that lead to resistance, a significant gap remains in our knowledge of resistance controlled by many loci and the evolutionary factors that influence the maintenance of resistance over time. Here, we perform a multi-level analysis involving whole genome sequencing and assembly, resequencing and gene expression analysis to both uncover putative loci involved in nontarget herbicide resistance and to examine evolutionary forces underlying the maintenance of resistance in natural populations. We found loci involved in herbicide detoxification, stress sensing, and alterations in the shikimate acid pathway to be under selection, and confirmed that detoxification is responsible for glyphosate resistance using a functional assay. Furthermore, we found interchromosomal linkage disequilibrium (ILD), most likely associated with epistatic selection, to influence NTSR loci found on separate chromosomes thus potentially mediating resistance through generations. Additionally, by combining the selection screen, differential expression and LD analysis, we identified fitness cost loci that are strongly linked to resistance alleles, indicating the role of genetic hitchhiking in maintaining the cost. Overall, our work strongly suggests that NTSR glyphosate resistance in *I. purpurea* is conferred by multiple genes which are maintained through generations *via* ILD, and that the fitness cost associated with resistance in this species is a by-product of genetic-hitchhiking.

## Introduction

Pesticide and herbicide use has reshaped ecological networks and induced strong selective pressures in the anthropogenic era. How species may adapt to strong selection is a fundamental question in evolution with great importance to the control of pesticide resistant organisms. A striking feature of pesticide resistance evolution is that there are a number of different genetic solutions that can lead to resistance^1,2^. In herbicide resistant plants, for example, resistance can be due to single gene mutations, often found in the herbicide’s target protein (target site resistance, TSR), or due to changes in multiple genes, often underlying nontarget herbicide resistance (NTSR) mechanisms^3,4^. A growing body of work has produced a better understanding of resistance controlled by single genes across a variety of species^3,5,6^. However, we currently lack a deep understanding of both the genetic basis and evolutionary potential of nontarget site resistance mechanisms genome wide^7–9^.

This is due in part to the broad nature of nontarget site herbicide resistance mechanisms more generally. NTSR can be caused by reduced herbicide uptake or penetration, altered translocation or sequestration, and/or herbicide detoxification^10–12^ -- mechanisms that likely rely on a complex genetic basis^8,13–15^. While some investigations have pinpointed a single gene conferring NTSR^16,17^, gene expression surveys or whole genome re-sequencing assays in a small handful of resistant weeds are beginning to shed light on the complexity of nontarget resistance mechanisms^18–20^. For example, in both *Amaranthus tuberculatus* and *Ipomoea purpurea*, a number of different loci found across the genome -- whether structural, regulatory, or both -- exhibit signs of selection and are thus putatively involved in resistance^18–20^. Because we lack a deep understanding of the genetic basis of NTSR in most weeds, however, we lack a firm grasp on the underlying forces that influence the maintenance of resistance in natural populations, such as the prevalence of alleles that may contribute to fitness costs of resistance, or the presence of interchromosomal linkage disequilibrium (ILD). The presence of ILD between unlinked regions of the genome would implicate the potential for epistatic interactions between alleles underlying either resistance or its cost.

*Ipomoea purpurea* is a common agricultural weed in the southeast and Midwest United States. Populations of this species, which have consistently been exposed to glyphosate based herbicides since the late 1990’s^21,22^, exhibit varying levels of herbicide resistance, with some populations exhibiting low and others high survival post-herbicide application^21^. There is a fitness cost associated with this resistance: resistant populations show lower germination and deteriorated seed quality compared to susceptible populations^23^. Further, populations from the south and midwest show evidence of genetic admixture, with both microsatellite and SNP data showing low genetic differentiation (F_ST_ = 0.11-0.14,^24^ and recent genetic connectivity^24^). RADseq and exome sequencing has identified regions of the genome under selection and thus associated with herbicide resistance. These regions are enriched for cytochrome P450s, glycosyltransferases, and ABC transporter genes, indicating a likely role of herbicide detoxification in conferring resistance^20^. Despite evidence that detoxification underlies resistance in this species, and suggestions that loci found on different chromosomes contribute to resistance, previous work relied on low-coverage RADseq sequencing without the benefit of a contiguously assembled genome. Thus, loci that may contribute to NTSR or its cost were likely missed^25^, meaning that we lack a thorough understanding of NTSR, the genomic context of NTSR alleles, and the potential for relationships among NTSR alleles in this species -- all crucial to understanding the evolution of resistance more broadly.

Here, we implemented a genome-wide selection screen using whole-genome resequencing of natural populations along with a gene expression survey to characterize the genetic architecture of glyphosate resistance and its cost in *Ipomoea purpurea*. We complemented our survey with a functional assay to test the potential that resistant *I. purpurea* individuals detoxify the herbicide. Given previous evidence that multiple loci likely contribute to herbicide resistance in this species, and evidence of fitness cost of resistance, we made two main predictions regarding genome-wide patterns of selection associated with resistance in *I. purpurea*. First, we expected that regions of the genome showing high differentiation and marks of selection when comparing herbicide resistant and susceptible individuals would contain loci with strong functional links to either herbicide resistance or its cost. Second, we anticipated that linkage disequilibrium, the non-random association of alleles at different loci, should be evident among regions of the genome housing resistance loci. Although inter-chromosomal linkage disequilibrium has been identified in other systems assessing ecologically relevant traits such as mate choice and coloration^26–28^, it is unknown if loci underlying herbicide resistance that are found across chromosomes exhibit long-distance or inter-chromosomal linkage disequilibrium, as would be expected if adaptation to herbicide is facilitated by multilocus genotypes favored by selection (*i*.*e*., coadapted gene complexes)^29–31^.

## Results

### A chromosome-scale genome assembly for common morning glory

We assembled a reference *I. purpurea* genome to test these hypotheses, generating the first genome sequence for this common and noxious weed. We generated a total of 48 gigabases of PacBio Sequel whole genome shotgun data (Supplemental Figure S1a). Based on a flow cytometry genome size (Benaroya Institute, Seattle, WA), this amounts to roughly 59X genome coverage for an estimated haploid genome size of 814 Mb. We used 34.79 gigabases of trimmed and self-corrected reads for assembly, scaffolding and polishing, which produced a 602 Mb assembly in 402 scaffolds (434 contigs), with a scaffold N50 of 5.77 Mb.

We performed pseudomolecule scaffolding with Phase Genomics Hi-C map, which collapsed the assembly into the expected 15 haploid chromosomes (Supplemental Figure S1b). We renamed and oriented chromosomes according to a high degree of synteny with the related *Ipomoea nil* genome^32^ (Supplemental Figure S1c). No misjoins were identified and broken based on the Hi-C linkage data. BUSCO scores on the unannotated assembly show 97.5% completeness against the Viridiplantae odb10 gene set (Supplemental Figure S1d).

Approximately 63% of the assembly was masked as repetitive DNA, with a significant proportion of recently-expanded Long Terminal Repeat (LTR) retrotransposons (Supplemental Figure S1e). Given the high degree of synteny with *I. nil* genome, the discrepancy between the flow cytometry genome size (814 Mb) and the assembled size (602 Mb) is likely due to young retrotransposon proliferation. We annotated 53,973 genes by combining ab initio gene predictions and RNA sequencing data from leaf tissue. The assembly shows a high degree of synteny with several genomes in the Convolvulaceae family, including *I. nil, I. trifida, and I. triloba (*Supplemental Figure S2).

### Detecting loci under selection

Whole-genome analysis of 69 individuals identified twenty-one regions across the genome exhibiting signals of strong genetic differentiation and signs of selection when comparing herbicide resistant and susceptible populations (Supplementary TableS4). These regions exhibited a G_ST_ and Md-rank-P in the top 5 percentile (G_ST_ > 0.284 and Md-rank-P > 5.69) with the Md-rank-P being a composite test of selection that incorporates nucleotide diversity, Tajima’s D, Fay and Wu’s H, and H12. This strategy identified 4.47 Mb of the genome showing signs of selection associated with herbicide resistance. The regions under selection were located across nine chromosomes, varied in size between 26kb-1272kb (Figure 1), and housed 358 genes, 202 of which could be functionally annotated (Supplementary TableS5).

**Figure 1.**
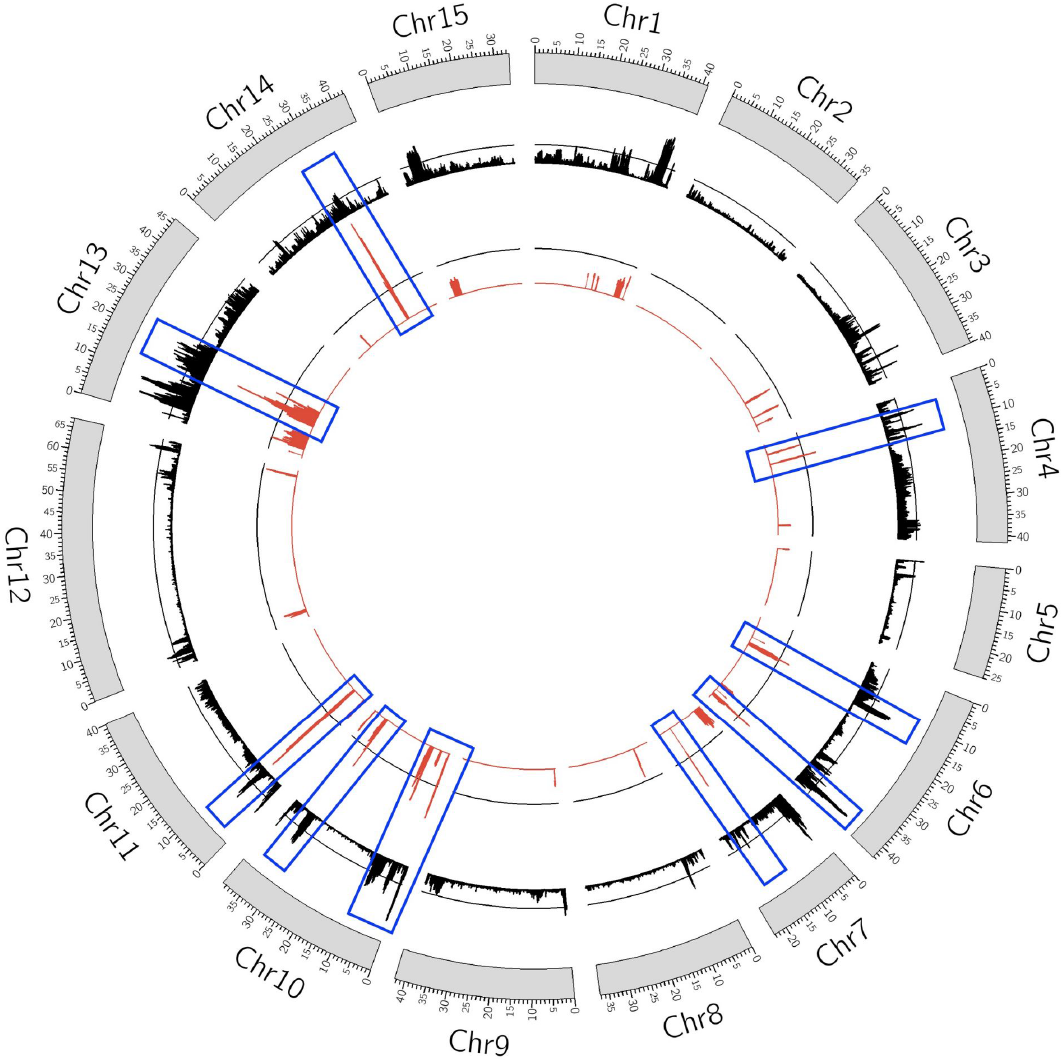
Circos plot depicting the regions of the genome that show signs of selection associated with herbicide resistance. The genome assembly resulted in 15 scaffolds which are represented here by grey bars. Values of G_ST_ describing the differentiation between the resistant and the susceptible populations are depicted by black bars, and the Md-rank-*P* values identifying signatures of selection are presented in red bars. Regions of the genome that exhibited both high differentiation (G_ST_ > 0.284) and a significant Md-rank-*P* value (Md-rank-P > 5.69) are identified by blue boxes. Black lines above both the G_ST_ and Md-rank-*P* represent 95% most extreme genome-wide values for each metric. Blue boxes on Chr5, Chr6, Chr10, Chr13, and Chr14 represent more than one region under selection.

The strongest signal of selection we uncovered was found within a 233kb region of chromosome 10 (average Md-rank-*P* = 7.99; average G_ST_ = 0.69, Figure 2). Within this region we identified 8 copies of cytochrome P450 genes (CYP) and 7 copies of glycosyltransferases, both of which are gene families previously implicated in herbicide detoxification. The eight cytochrome P450s belong to the 76A family (three CYP76A1 and five CYP76A2) and were present in tandem within 53kb. Four copies of the cytochrome P450s exhibited multiple non-synonymous mutations that were almost fixed in the resistant individuals (allele frequency = 0.95). Further, two of the eight cytochrome P450s (CYP76A2) in this block exhibited either a premature stop codon and/or a splice site donor variant (G->C) in the first intron (allele1 susceptible frequency = 0.68, resistant frequency = 0.05) in the majority of the susceptible individuals. The seven glycosyltransferases were found in tandem; one glycosyltransferase copy showed the loss of a stop codon (susceptible frequency = 0.64, resistance frequency = 0.05), whereas the other glycosyltransferases exhibited multiple non-synonymous mutations close to fixation in the resistant individuals (resistant frequency = 0.05, susceptible frequency = 0.60; Supplementary Figure S3). Additionally, the block of glycosyltransferases in this region showed evidence of a hard sweep (glycosyltransferases H12 = 0.87).

**Figure 2.**
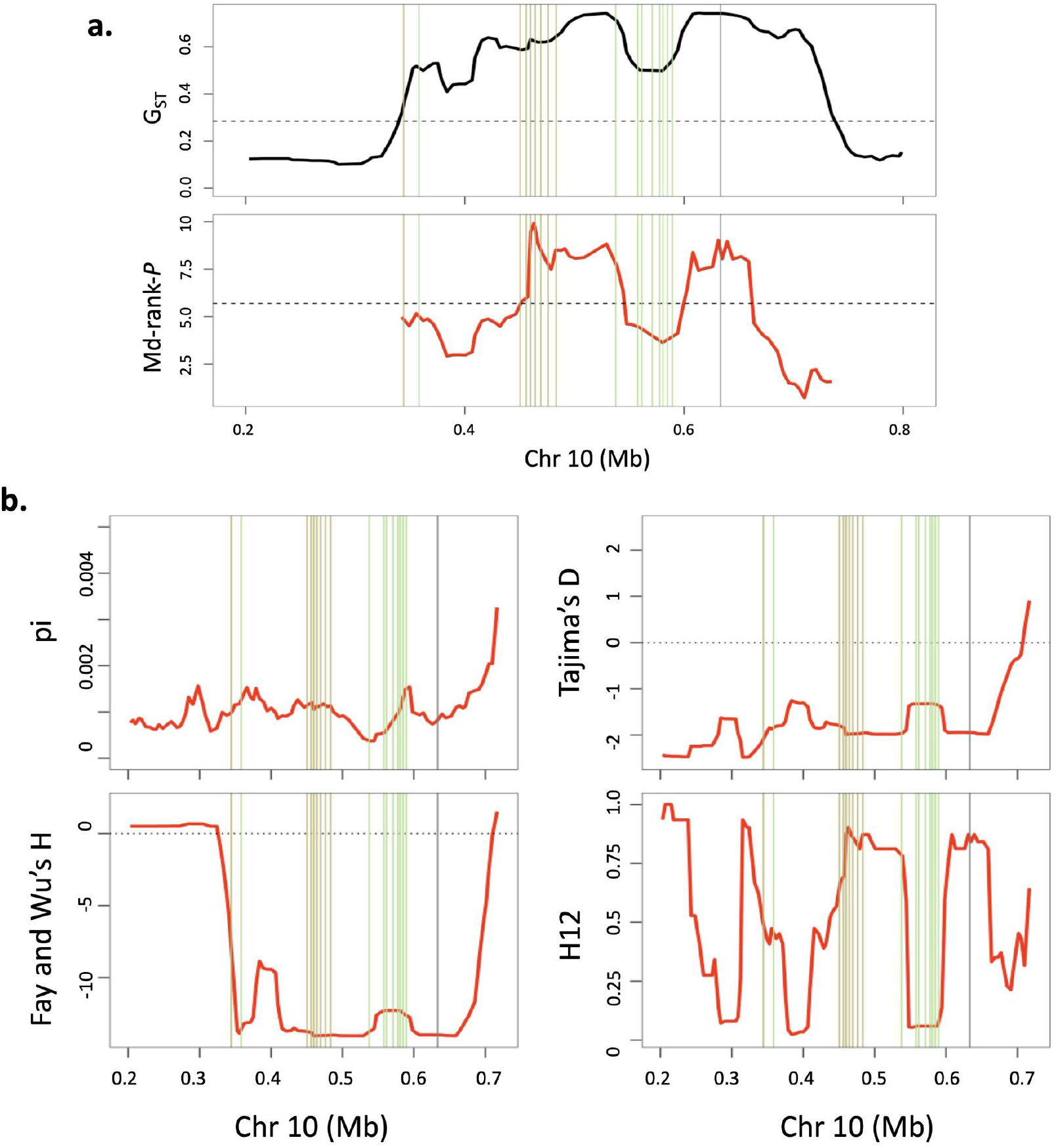
Region of Chromosome 10 showing signs of selection. Shown is the (a) G_ST_ (upper) and Md-rank-*P* (lower) for the resistant individuals which was estimated using statistics shown in (b) clockwise starting from upper left, pi, Tajima’s D, H12 and Fay and Wu’s H. Red lines indicate respective values for the resistant populations. Khaki vertical lines represent copies of glycosyltransferases, green vertical lines are the cytochrome P450, and the grey vertical line represents CTR1 (see below). The black dashed line in (a) represents 95 percentile values.

On chromosome 11, we found a 6.4-7.1Mb region to show signs of selection (average Md-rank-*P* = 8.91; average G_ST_ = 0.392) with six copies of a phosphate transporter gene (*PHO1*), an ABC transporter gene (*ABCB19*), and a sugar transporter gene (*ERD6*), all containing almost fixed non-synonymous mutations in resistant individuals (resistant frequency = 0.99). This region also contained two copies each of a glycosyltransferase and a cytochrome P450 gene (CYP736A12 family, Supplementary TableS5).

Another region of note showing strong signals of selection was found on chromosome 6 (average Md-rank-*P* = 6.55; average G_ST_ = 0.782, Figure 1), with evidence of strong differentiation continuing further upstream and downstream (40.23Mb - 40.81Mb; mean G_ST_ = 0.727). Within the extended downstream region, we found ethylene responsive transcription factor (*ERF4*) and multiple copies of serine/threonine kinases, genes that are involved in the signal transduction in response to various biotic and abiotic stresses^33–37^. Within this region we also identified loci that are likely related to the cost of resistance in this species, expanded upon further in ‘Signs of selection on potential cost loci’ below.

Across the other regions exhibiting signs of selection, we found multiple environmental stress response genes (Supplemental TableS4): serine/threonine-protein kinase CTR1 (chr10) involved in stress signalling, *LOG3* (chr 4), associated with drought stress response^38^, the GT-3B transcription factor (chr 6) which is responsible for inducing response to salt^39^, tubby-like F-box protein 5 (TULP5), AP2-like ethylene-responsive transcription factor PLT1, AT-hook motif nuclear-localized protein 24 (AHL24)^40^, and several homologs of FRS related sequence (FRS) and E3 ubiquitin-protein ligases, genes involved in response to oxidative stress^34,37^. Of special note, we uncovered a gene involved in the shikimate acid pathway on chromosome 10 -- the bifunctional 3-dehydroquinate dehydratase/shikimate dehydrogenase (DHD/SHD) gene, which is responsible for converting dehydroquinate to shikimate^41^. This latter gene is notable in that it is a part of the shikimic acid pathway, which is the biochemical pathway inhibited by glyphosate^42^.

Overall, our selection screen using a WGS resequencing approach identified highly differentiated regions under selection, with these regions containing genes involved in herbicide detoxification (cytochromeP450s, glycosyltransferases, ABC and phosphate transporters), the shikimate acid pathway (DHD/SHD), environmental sensing (serine/threonine kinases), and stress response genes (ERFs, PLT1, E3 ubiquitin-protein ligase, FRSs, TULP5, AHL24, LOG3, GT-3B transcription factor). Thus, our study expands on our previous work which found detoxification genes to be under selection^20^ by providing strong evidence that glyphosate resistance in *I. purpurea* is controlled by a polygenic NTSR mechanism likely involving herbicide detoxification, response to environmental stimuli and stress, and components of the shikimate acid pathway, which is the biochemical pathway inhibited by glyphosate.

### Gene expression differences implicate herbicide detoxification

We compared gene expression between herbicide treated resistant and susceptible plants and found support for the idea that herbicide detoxification, plant signalling, and stress response underlies resistance. Of the 250 differentially expressed genes (111 upregulated and 139 downregulated; Supplementary TableS6), we found cytochrome P450s, glycosyltransferases, and glutathione S-transferase genes (Figure 3a) to be differentially regulated between resistant and susceptible plants. Two copies of the cytochrome P450 family CYP82D7 were significantly upregulated in the resistant individuals (logFC: 2.05 and 1.35), along with two copies of UDP-glycosyltransferases (UGT87A2 and UGT88B1) and a glutathione S-transferase (GST). We additionally found a cytochrome P450 (CYP82C4) and a glycosyltransferase (UGT89B2) downregulated in the resistant individuals.

**Figure 3.**
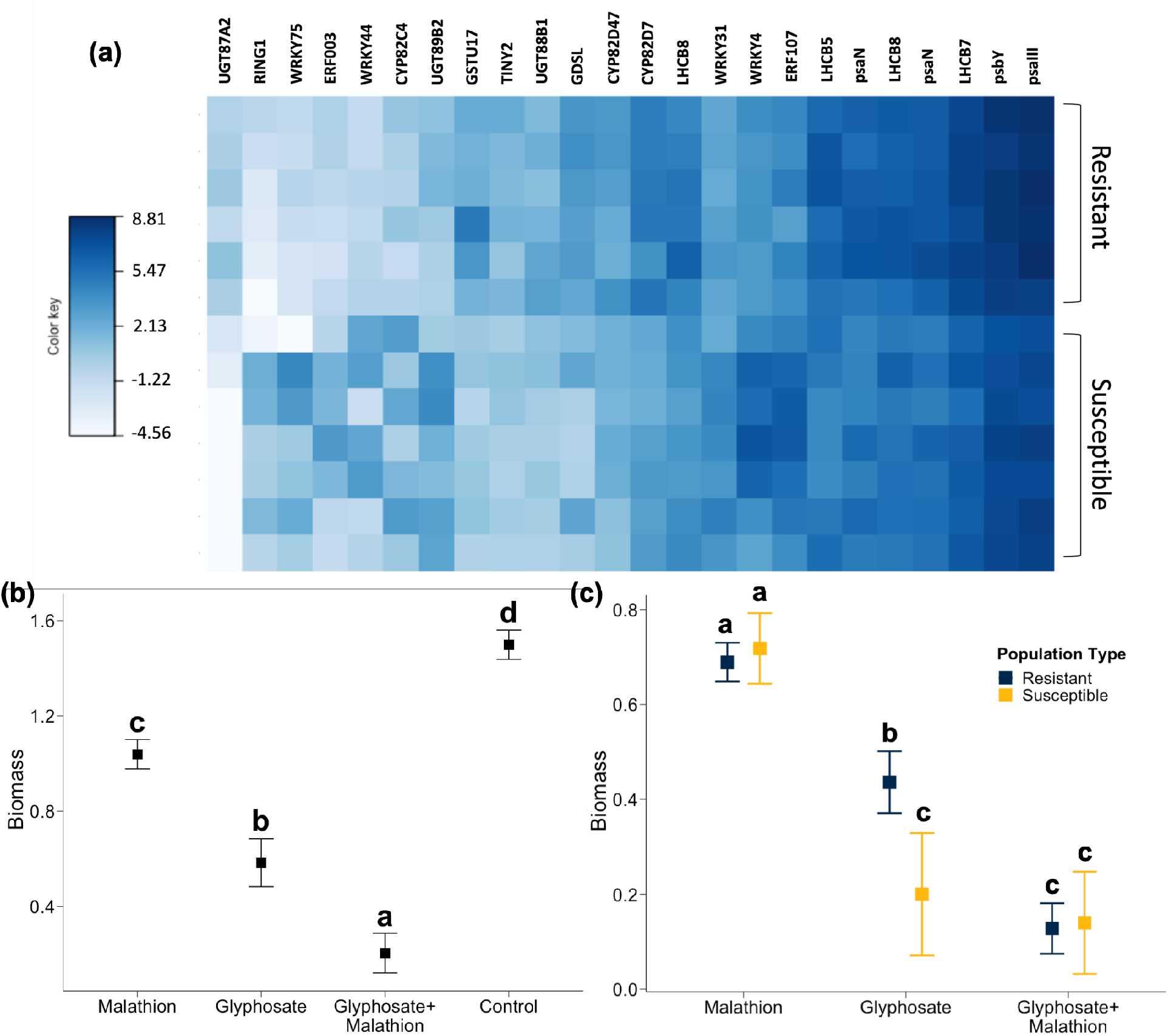
Gene expression variation associated with herbicide resistance, and results of a functional assay supporting the idea that resistance in *I. purpurea* is due to detoxification. (a) Loci associated with glyphosate resistance identified by differential expression analysis with P-value < 0.0005. Color key represents log2 fold-change values. (b) Least square means of above-ground biomass according to treatment (malathion, glyphosate, glyphosate plus malathion, and a control (no treatment)) and (c) summarized according to resistance type (R/S). Letters in (b) and (c) indicate significant differences between treatment environments. The addition of the cytochrome P450 inhibitor malathion reverses glyphosate resistance (glyphosate vs glyphosate+malathion, contrast estimate = 0.379, t-ratio = 2.946, p-value = 0.019), with the resistant individuals showing the same phenotype as the susceptible individuals in the presence of glyphosate and malathion but not in the presence of glyphosate only.

We likewise uncovered differences in the expression of genes associated with environmental stress responses. Among notable genes were ethylene responsive transcription factors (ERF003, ERF107, TINY^43,44^), serine/threonine kinase BLUS1^45^, E3 Ubiquitin protein ligase PUB23^34^, NAC domain containing protein 72^46^, and WRKY transcription factors (WRKY4, WRKY31, WRKY75^47–49^) (Figure 3a). Homologs of these genes (ERF4, PLT1, CTR1, PRP4, HT1, B120, RHC1A, RF298, NAC92, NAC25, WRKY22; Supplementary TableS5) were also under selection when comparing herbicide resistant and susceptible populations.

In the control (non-herbicide) environment, we found 623 differentially expressed genes when comparing resistant and susceptible individuals (319 upregulated and 304 downregulated; Supplementary TableS7). We identified multiple copies of cytochrome P450s, glycosyltransferases, and ABC transporters that were differentially expressed, indicating that glyphosate resistance through detoxification is constitutive, and not induced, in this species. Interestingly, the specific cytochrome P450s and glycosyltransferase genes that exhibited signs of selection from our whole-genome scan were not the same as those that exhibited differential expression, which could be due to the non-simultaneous nature of the gene transcription response to glyphosate^50^, or could represent a transcriptional sampling stage caveat.

### Functional assay supports herbicide detoxification as a mechanism of resistance

We performed an assay to determine if the functional mechanism of resistance in *I. purpurea* was herbicide detoxification (following^17,51–54^). We applied malathion, a pesticide that inhibits cytochrome P450s, to multiple resistant and susceptible *I. purpurea* individuals from the same populations used in the WGS re-sequencing and gene expression studies. The expectation that malathion would act to inhibit *I. purpurea* cytochrome P450s was met; we found a significant overall treatment effect (F-value = 59.33, df = 3, p < 0.0001; Fig 3b) with individuals treated with both glyphosate and malathion showing lower biomass compared individuals treated with either malathion, glyphosate, or untreated controls (Fig 3b; Supplemental TableS8).

As expected, resistant individuals showed significantly greater biomass compared to the susceptible individuals in the presence of glyphosate (F-value = 4.81, df = 1, P-value = 0.03; Figure 3c). However, the biomass of resistant individuals in the presence of both malathion and glyphosate was significantly lower than that of resistant individuals treated only with glyphosate (resistant plants, malathion+glyphosate *vs* glyphosate: t = 3.65, df = 78, p-value = 0.001), indicating that the presence of malathion reduces the resistance response. In fact, the presence of both malathion and glyphosate led to similar (and low) remaining biomass of both resistant and susceptible individuals (malathion+glyphosate treatment: resistant vs susceptible plants: t = 0.15, df = 32, p-value = 0.88). This shows that the presence of a cytochrome P450 inhibitor lowers the level of glyphosate resistance in *I. purpurea* plants, supporting the idea that modification to the detoxification pathway underlies glyphosate resistance in this species.

### Role of long-distance and interchromosomal linkage disequilibrium in maintaining NTSR alleles

Our whole-genome scan identified regions under selection containing genes involved in environmental sensing, stress responses, and herbicide detoxification. This broad scan implicates a polygenic basis of resistance in *I. purpurea* and shows that multiple regions of the genome likely contribute to resistance. We thus sought to determine if there was evidence of linkage disequilibrium between these regions, which would potentially suggest either epistatic interactions among alleles or the inheritance of coadapted gene complexes^29,55^. We calculated a measure of linkage disequilibrium (r^2^) between long-distance and interchromosomal SNPs that showed the most extreme level of differentiation and selection (98th percentile, G_ST_ > 0.39) -- regions on Chromosome 4, 6 (two regions, hereon referred to as 6.1 and 6.2), 10 and 11, and compared it to the whole-genome measure. We found that the five regions under selection showed islands of elevated interchromosomal linkage disequilibrium (Supplementary TableS9) in a backdrop of nearly zero genome-wide ILD (background interchromosomal r^2^ mean = 0.00096; Figure 4). Additionally, the five regions with high differentiation under selection also showed higher linkage (99th percentile ILD = 0.23 ± 0.0004 SE) in comparison to the five random highly differentiated regions of the same size that are not under selection (99th percentile ILD = 0.13 ± 0.0003 SE). The region under selection on Chr10 exhibited the strongest linkage to other chromosomal regions under selection (99% ILD Chr4-Chr10 = 0.256, Chr6.1-Chr10 = 0.257, Chr6.2-Chr10 = 0.22, Chr11-Chr10 = 0.17).

**Figure 4.**
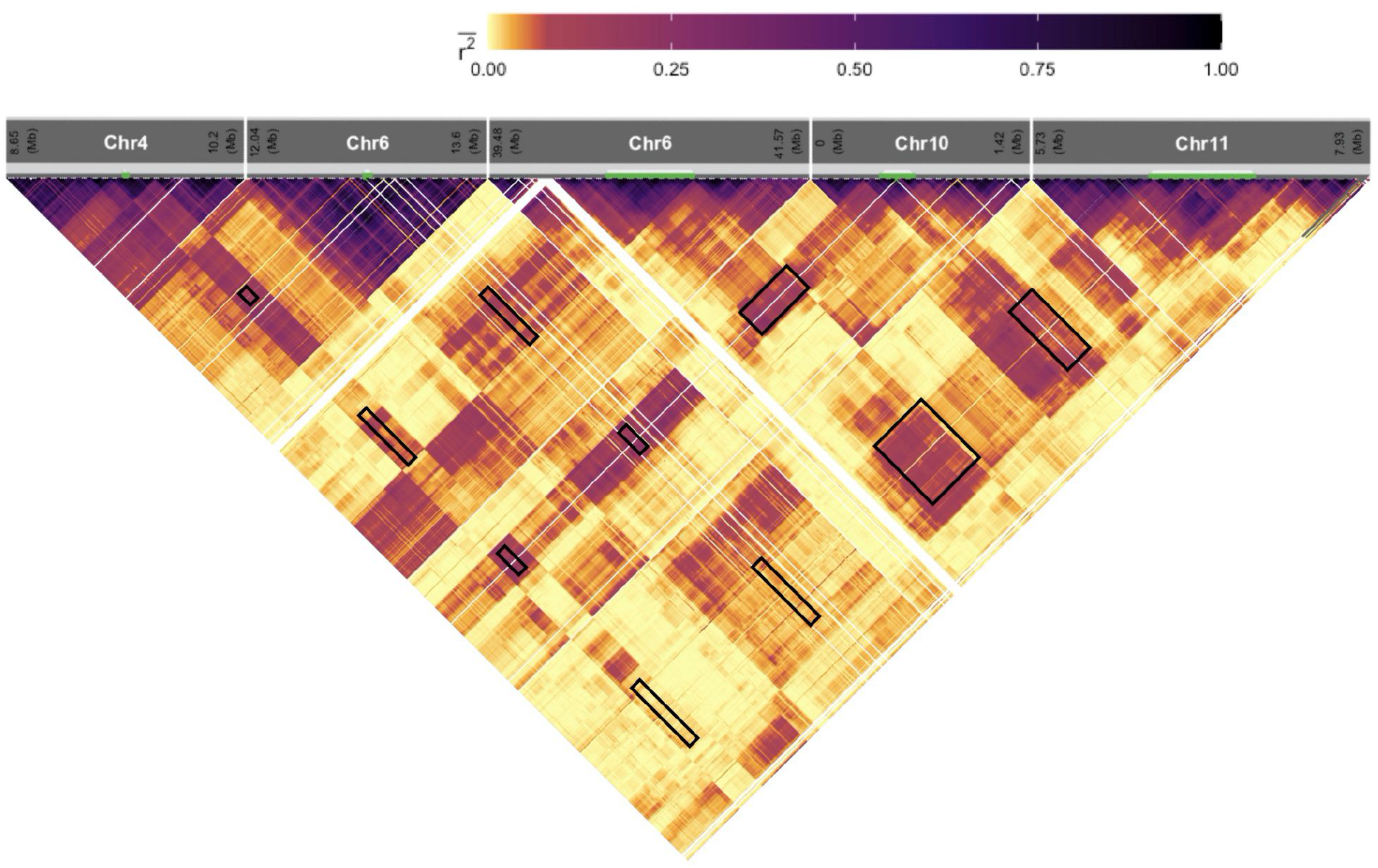
Long-distance linkage disequilibrium and ILD among the five highly differentiated (G_ST_ > 0.39) regions under selection associated with glyphosate resistance. The five intervals displayed islands of increased linkage disequilibrium as estimated by r^2^ for SNPs separated by at least 1 kb in and between broad regions under selection. The white lines represent absence of SNPs (missing data) whereas the black boxes represent linkage between the five selection intervals. r^2^ values are averaged over two-dimensional bins of 10 × 10 kb.

Interestingly, the highest r^2^ values (within the top 1 percentile) within these regions was observed for putative resistance genes identified above. For instance, multiple glycosyltransferases and cytochrome P450s under selection on Chr10 showed high ILD with SNPs on Chr11 (Supplementary TableS10). Multiple cytochrome P450 genes (*CYP76A2*) on Chr10 showed a high value of ILD with an uncharacterized protein and a region upstream of GT-3B on Chr6.1 (range of r^2^ = 0.256-0.278, Supplementary TableS10) as well as the intergenic region between the transcription factors *SPL1* and *DOF1*.*4*, both of which are responsible for plant growth and development, on Chr6.2 (range of r^2^ = 0.249-0.288), perhaps indicating that this region on Chr6 may influence the regulation of the *CYP76A2* on Chr10 in some way. Furthermore, the highest linkage between Chr4 and other chromosomes (range of r^2^ = 0.167-0.274) was observed for a SNP just upstream of *LOG3* on Chr4 (Supplementary TableS9). Thus, the identified resistance alleles within these five highly differentiated regions show signs of linkage and perhaps evidence of epistatic selection.

Local regions of strong long-distance linkage disequilibrium and ILD within species might be aided by demographic processes like population structure^55,56^, genetic drift^57^, or could be due to other processes like selection^28,58^. Furthermore, epistatic interactions among loci wherein adaptive alleles at two independent loci will be inherited together can generate linkage disequilibrium^28,58–60^. Given that our sampling design included multiple resistant and susceptible populations from varied locations, low population differentiation among populations, and evidence of recent migration between them (Supplementary Figure S4), it is unlikely that the observed ILD is due entirely to demographic processes. Moreover, we observed the strongest ILD between regions under selection harboring resistance associated genes, indicating the potential role of selection in maintaining the observed ILD. Thus, our finding suggests that the highly differentiated regions under selection containing candidate loci for glyphosate resistance are linked, indicating the potential role of epistatic selection in maintaining resistance.

### Signs of selection on potential cost loci

Our scan of regions associated with herbicide resistance, paired with a transcriptome survey, identified potential alleles with strong functional connections to the previously identified fitness cost in resistant *I. purpurea*. We found alternate alleles close to fixation in each population type within the 585kb highly differentiated region on chromosome 6 (40.23Mb - 40.81Mb; mean G_ST_ = 0.727, Figure 5a). This region contained the nuclear fission defective 6 (*NFD6*) and NAC transcription factor 25 (*NAC25*) genes, both of which function in seed development. *NAC25*, a gene that is required for normal seed development and morphology ^61^, exhibited two missense variants in the resistant individuals (mutant allele resistant frequency = 0.91, susceptible frequency = 0.23). Additionally, *NFD6*, a protein required for nuclear fusion in the embryo sac during the production of the female gametophyte^62^, contained six missense variants in the resistant individuals (mutant allele resistant frequency = 0.88, susceptible frequency = 0.21). The resistant haplotype of this gene also contained 10 SNPs in the promoter region which could potentially alter its expression. Indeed, we found this protein to be downregulated in the presence of the herbicide, with a log-fold change of -3.52 in resistant individuals as compared to the susceptible individuals (Figure 5b). Thus, our data suggest that these genes may be responsible for the lower and abnormal germination leading to the observed fitness cost in this species ^23^).

**Figure 5.**
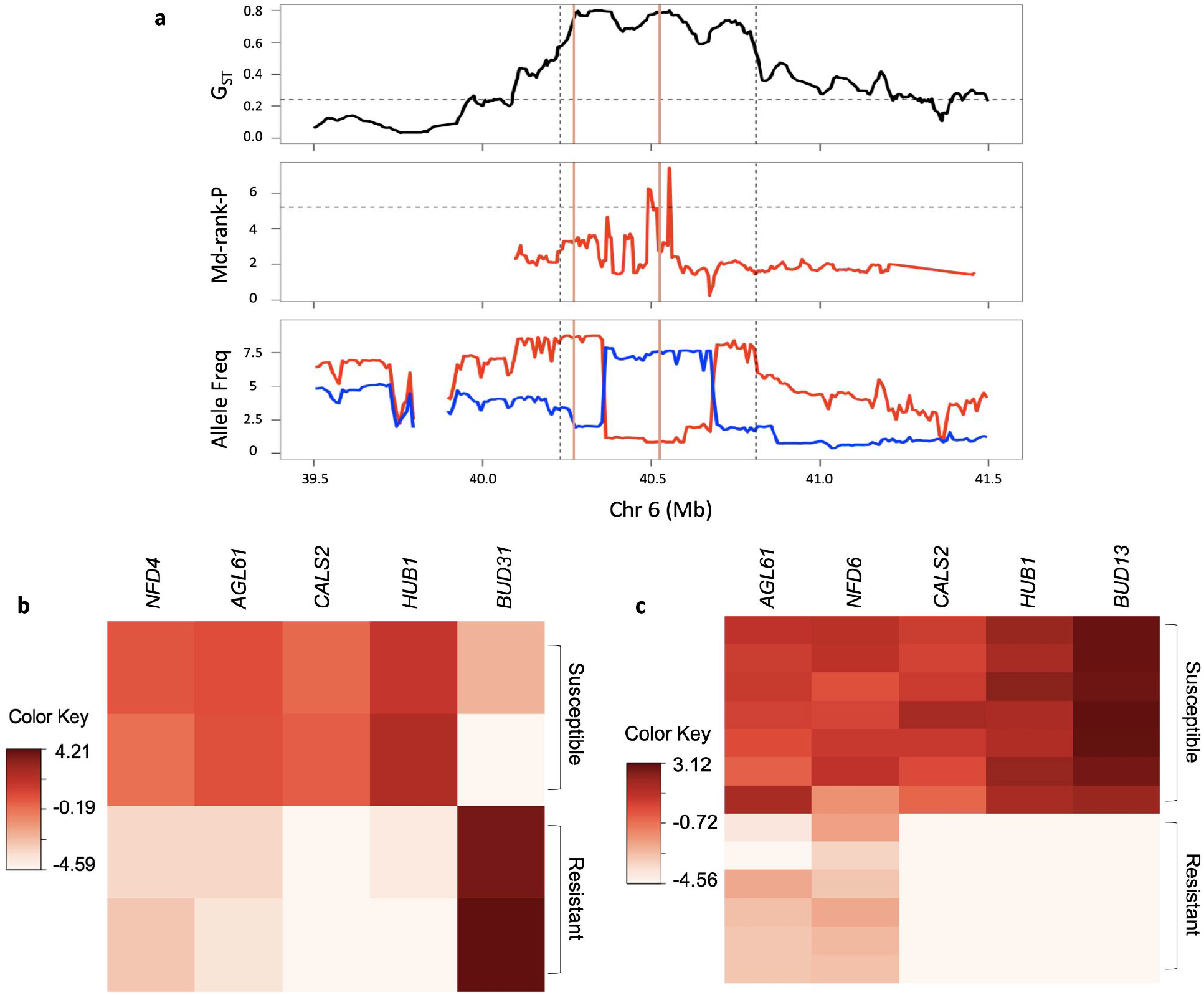
Loci associated with the cost of glyphosate resistance identified by the (a) whole-genome selection-scan and differential expression analysis in the (b) absence and (c) presence of herbicide. Top panel of (a) represents G_ST_ between the resistant and the susceptible populations, mid panel is the Md-rank-*P* value, and the lower panel represents the allele frequency. Salmon vertical lines represent NFD6 and NAC25, in that order. Red and blue represent resistant and susceptible populations, respectively. Black horizontal dotted lines represent 95 percentile values while vertical lines represent regions with G_ST_ above 0.6. The differentially expressed cost genes shown here were chosen based on their functional annotation and had FDR < 0.005 and P-value < 0.00005. Color key represents log2 fold-change values.

Interestingly, this highly differentiated region containing these seed development genes is strongly linked to other regions under selection that harbor resistance alleles (99% ILD value = 0.22; Figure 4), indicating the potential role of linkage disequilibrium in maintaining the cost. More specifically, *NFD6* is within 83 kb from, and thus physically linked to (r^2^ = 0.70), the potential regulatory region on Chr6.2 that exhibits interchromosomal long distance linkage disequilibrium with the CYP76A2 gene on Chr10 (ILD = 0.27). Further, the NAC25 gene is found in close proximity to serine/threonine kinases on Chr6.2 (i.e., 82 kb away), indicating another potential gene involved in the cost phenotype is in physical linkage (r^2^ = 0.67) with potential NTSR loci.

In addition to the potential cost loci identified from the WGS screen, our gene expression analyses in the absence of herbicide (*i*.*e*. the environment in which fitness costs are assessed) comparing resistant and susceptible individuals identified five differentially expressed gene that play a role in fertilization and seed maturation and are thus potentially related to the cost (Supplementary TableS6). Of special interest, the bud-site selection protein 31 (*BUD31*) was found to be highly upregulated (logFC = 7.22) in resistant plants in the absence of herbicide, whereas its homologue *BUD13* was highly significantly downregulated in resistant individuals in the presence of herbicide (logFC = -11.39). *BUD13* is involved in pre-mRNA splicing in embryos and is critical for early embryo development^63^.

In the control environment, two genes downregulated in the resistant individuals -- *NFD4* (logFC = -3.61) and Agamous-like MADS-box protein AGL61 (*AGL61*; logFC = -4.98) -- are involved in megagametogenesis. The *NFD4* gene, like *NFD6*, is responsible for ovule polar nuclei fusion during female karyogamy^62^, whereas *AGL61* is required for the central cell development and differentiation^64^. A loss of function mutation in *AGL61* has been shown to cause abnormal morphology and over 50% seed abortion upon fertilization in Arabidopsis^64^. We also identified a callose synthase 2 (*CALS2*) to be strongly downregulated in resistant individuals (logFC = -8.72); another member of the callose synthase family (*CALS5*) has been shown to be responsible for pollen viability^65^. Finally, we also found that E3 ubiquitin-protein ligase BRE1 (*HUB1*), a protein involved in seed germination, was strongly downregulated among the resistant individuals (logFC = -8.27). *HUB1* has been shown to control chromatin remodeling during seed development and leads to alterations in seed dormancy^66^.

Interestingly, three of these five candidate genes (*AGL61, CALS2, HUB1*), and homologues of other two (*BUD13* and *NFD6)* were also significantly downregulated in the resistant populations in the presence of herbicide (Supplementary TableS6). These candidate cost genes are all essential for plant reproduction and are highly downregulated (except *BUD31*) in the resistant population in both the absence and presence of the herbicide, and thus could potentially explain the phenotypic costs of glyphosate resistance in *I. purpurea* seen by Van Etten and colleagues^23^.

## Discussion

While there is an increasing appreciation for the role of nontarget site mechanisms underlying herbicide resistance in agricultural weeds^12,67–69^, there are strikingly few comprehensive whole genome assays of resistant weeds suggesting that the entirety of the NTSR response is rarely captured. Our study using a sequenced and assembled genome, whole genome resequencing of natural populations, and a gene expression survey offers a unique opportunity to identify loci associated with NTSR and to further investigate the evolutionary forces that underlie the maintenance and expected evolutionary trajectory of resistance alleles in natural populations.

Our results show detoxification underlies resistance in *I. purpurea*. Detoxification is hypothesized to involve three steps -- uptake of the herbicide by phosphate transporters, chemical modification (*i*.*e*. the addition of an OH and sugar group by cytochrome P450s and glycosyltransferases, respectively), and transport to vacuoles by ABC transporters and other sugar transporters where the molecule is stored and/or inactivated^12,18^. We found evidence of selection on all genes involved in this pathway. We also found evidence of selection (and in some cases, differential expression) of genes involved in plant signalling and environmental stress (i.e., serine/threonine kinases, ERFs, E3-Ubiquitin Ligases, FRSs, LOG3, GT-3B, TULP5) as well as genes involved in the shikimate acid pathway (DHD/SHD). Our results thus expand what we currently know about the detoxification NTSR mechanism in this species to include plant signalling and stress responses, both of which are either hypothesized ^7^ or shown to be involved in herbicide resistance^70–73^. While we do not currently have functional genomics resources for this species, our study using a cytochrome P450 inhibitor verifies that resistant *I. purpurea* individuals have the ability to detoxify the herbicide. The next step in understanding resistance in *I. purpurea* involves determining the contribution of each of the candidate loci under selection (and/or showing differential regulation) to both resistance and its associated cost. With future development of genome editing protocols for *I. purpurea*, we will be able to experimentally test the function of loci hypothesized to be contributing to herbicide resistance.

Due to the involvement of multiple genes involved in the herbicide detoxification pathway, and evidence for selection on regions of the genome found on separate chromosomes, we hypothesized that multiple loci would show evidence of ILD, perhaps indicating epistatic selection between alleles or co-inheritance. Our results support this hypothesis. Foremost, in contrast to low background ILD, long-distance linkage disequilibrium and ILD were high among intervals under selection, and consistently differentiated between the resistant and susceptible types across multiple populations. The strongest linkage was observed between putative resistance genes that exhibited signs of selection. This linkage could quickly become very steep in the presence of epistatic interactions among loci^74^, as would be the case if genes underlying NTSR worked in concert to produce the resistance phenotype. Indeed, we found high ILD values between regulatory regions and resistance alleles, and between intervals harboring genes involved in the same molecular pathways (e.g. detoxification, and stress signaling and response).

Although linkage should become decoupled over time due to recombination and gene flow, the ongoing selection for herbicide resistance could slow down this decoupling between these functionally interacting genes^75^. Even given gene flow between these populations ^24^, co-adaptation and epistasis could lead to the fixation of the resistance alleles given strong selection^76^ or weaker recombination rate ^77^. Thus, long-distance linkage disequilibrium and ILD aided with epistatic selection could act to maintain resistance through generations in natural populations.

One evolutionary force that should counteract the continued evolution of resistance is the potential for fitness costs of resistance, either due to the pleiotropic effects of resistance alleles themselves or due to negative fitness effects of loci that are linked to resistance loci. While costs are central to theories of resistance evolution^78–81^, there are currently no examples, to our knowledge, in which the loci underlying fitness costs of nontarget site resistance have been identified. Our results suggest candidate loci associated with the previously identified cost of glyphosate resistance. Specifically, we found a highly differentiated region on Chr6 that exhibited alternate alleles in resistant and susceptible populations, and found this region to contain loci required for normal seed development and maturation (*NAC25, NFD6*). One of these genes, *NFD6*, was differentially regulated in the resistant individuals, further supporting its role in the low seed quality, and thus fitness cost, that we have previously described^23^.

Additionally, our results strongly suggest genetic hitchhiking may act to maintain the cost in this species. Both *NAC25* and *NFD6* are physically linked on chromosome 6 to the regions under selection containing serine-threonine kinase genes and a regulatory region that is itself exhibiting ILD to the *CYP76A2* gene on chromosome 10. Although recombination should decouple cost alleles that are physically linked to resistance alleles, these loci would not completely decouple if the recombination rate (c) is much lower than the selection coefficient (s) (i.e., c<<s,^82^. The requirement that c<<s is not improbable given the close proximity of cost and resistance loci (< 85kb) and the strong ongoing selection for herbicide resistance. Furthermore, if the ratio c/s < 10^−4^, the hitchhiking would almost be complete and the cost alleles could become fixed in the populations^83^. Alternatively, it is possible that new compensatory mutations arising in the population could increase in frequency over time due to selection, decoupling the cost and resistance alleles and thus reducing fitness cost associated with glyphosate resistance^84,85^.

Overall, our work identified the potential genetic basis of NTSR glyphosate resistance in *I. purpurea* -- our whole genome and transcriptome assays strongly support the role of detoxification conferring herbicide resistance in this species, and we additionally identified a role for plant sensing and stress along with components of the shikimate acid pathway. Interestingly, we show that NTSR glyphosate resistance in *I. purpurea* is conferred by multiple loci which are maintained through generations via ILD. We also provide strong evidence to support the idea that fitness costs may be due to loci in strong linkage with resistance loci. Our work highlights the importance of multi-level, multi-population study in identifying the genetic mechanisms underlying polygenic defense traits, and for understanding the complex genetic-interplay between defense and cost.

## Methods

### Genome sequencing, assembly, and annotation

We used an *I. purpurea* line originally sampled from an agricultural field in Orange County, NC, in 1985 by M. Rausher (*i*.*e*., prior to the widespread use of glyphosate) and selfed for >18 generations in the lab for genome sequencing (seeds of this line ‘Fred/C’ are available upon request). High molecular weight DNA was isolated from flash-frozen leaf tissue using a modified large-volume CTAB protocol^86^ and sequenced on a PacBio Sequel at the University of Georgia. Raw PacBio subreads from 9 cells of Sequel chemistry were error-corrected with Canu (v1.7.1)^87^ with default parameters for raw PacBio reads (--pacbio-raw). The corrected and trimmed reads from Canu were assembled with Flye (v2.4-release)^88^ and anchored onto pseudomolecules by nearly 81 million read pairs of Phase Genomics Hi-C (Seattle, WA) of leaf tissue using Sau3AI cutsites. Within-genome and across-genome synteny was visualized using the CoGE SynMap platform^89^, with DAGChainer options “-D20 -A 5”, as well as with jcvi with default parameters (https://github.com/tanghaibao/jcvi). *Ipomoea purpurea* pseudomolecules were numbered and oriented according to chromosome synteny against *Ipomoea nil* pseudomolecules (Supplemental Figure S2).

Raw 50nt single-end RNA-seq reads were aligned using STAR (v.2.7.0)^90^ with default single-pass parameters. Repetitive elements were first annotated with RepeatModeler (v1.0.11). Long Terminal Repeat (LTR) retrotransposons were annotated with LTRharvest (v1.6.1) with options -similar 85 -mindistltr 1000 -maxdistltr 15000 - mintsd 5 -maxtsd 20”. RepeatModeler annotations were combined with all Viridiplantae repeats from Repbase and used as a species-specific repeat database built using RepeatModeler with default options.

Genome annotation was performed using a diverse set of evidence. First, a set of 12 RNA-seq libraries from leaf tissue was aligned with STAR (v2.7.0), and transcripts assembled with Stringtie (v2.1.3). MAKER2^91^ was initially run with evidence from the RNA-seq alignments, as well as peptides from *I. trifida, I. triloba*, and *I. nil*. The resulting gene set was used to train SNAP (v2013-11-29). AUGUSTUS (v3.3.2) was trained with evidence from BUSCO (v4.1.0) against the eudicot odb10 set. with default options. MAKER2 was re-run with the *ab initio* SNAP and AUGUSTUS training sets, in addition to the homologous protein and RNA-seq evidence, to build a final gene annotation set.

### Sampling and sequencing

We selected eight populations to investigate the genetic basis of glyphosate resistance and its cost following^20^-- 4 low resistance, from here on referred to as the susceptible population (S: <20% population survival at 1X the field dose of RoundUp) and 4 high resistance populations (R: >70% population survival at 1X the field dose of RoundUp), from here on referred to as the resistant populations (Supplementary TableS1). Seeds from 10 maternal lines per population were germinated, except for one susceptible population (RB), wherein 9 maternal lines were used. We extracted DNA from leaf tissue using the Qiagen Plant DNeasy kit. 150 paired-end sequencing was performed using Illumina HiSeq4000 and NovaSeq6000 using three and two lanes, respectively. We sequenced two populations at high coverage (at least 25X) and the remaining six populations at low coverage (10X). Two populations (WG, resistant and RB, susceptible) were run on one lane of HiSeq6000 and NovaSeq6000 each whereas the other lane had the remaining six populations. This yielded a total of 3,300,397,148,700 bases with average coverage of 28.84X for WG and RB. Coverage of the other six populations has an average of 14.66X.

### Variant calling

We aligned the reads to our draft genome using BWA mem v0.7.15^92^ with parameter -M. Since the same sample was sequenced using multiple platforms (HiSeq and NovaSeq), the alignment files were merged and duplicate reads were marked using the MarkDuplicate tool of Picard v2.8.1 (http://broadinstitute.github.io/picard). Next, we prepared a database of true known variants, required for base recalibration. This database was created using data from the top eighteen individuals with the highest read counts, upon which variant call was performed using the HaplotypeCaller tool of GATK v4.1^93^. Low confidence variants were filtered out using the VariantFiltration tool of GATK v4.1^93^ (15 < DP < 60; ReadPosRankSum < -8.0; QD < 2.0; FS > 60.0; SOR > 3.0; MQ < 40.0; MQRankSum < -12.5) and only the high confidence variants were used in the dataset. This was used to recalibrate base qualities using GATK v4.1 tools BaseRecalibrator and ApplyBQSR^93^. Variants were called individually on all the individuals using the HaplotypeCaller tool of GATK v4.1^93^ using parameters -ERC GVCF --min-pruning 1 --min-dangling-branch-length 1. The variants from each individual were combined to one variant file (a raw cohort variant file) using the tools GenomicsDBImport, GenotypeGVCFs, and GatherVcfs^93^, with invariants included. Next, multiple rounds of filtration were performed on this variant dataset to filter out potential false positives. First, using the GATK v4.1 tools VariantFiltration and SelectVariants we filtered the variants using the parameters QD<1.5, DP<10 and DP>2000, FS>80, SOR>5, MQ<40, MQRankSum< -6 and MQRankSum>6, and ReadPosRankSum< -4 and ReadPosRankSum> 4^93^. For the next round of filtration, we removed variants that had genotype depth more than twice the average and heterozygosity more than 0.8 using the het packages from VCFtools v0.1.15^94^. In the third round of filtration, we filtered variants that had quality above 20, had no missing information, a minor allele frequency of 0.05, and a minimum mean depth of 10 (vcftools --minQ 20 --max-missing 1.0 --maf 0.05 --min-meanDP 10)^94^. Finally, we filtered using BCFtools (v1.7)^95^ to keep only bi-allelic SNPs (bcftools view -m2 -M2 -v snps). This gave us a total of 3,942,549 high confidence SNPs. These SNPs were used for downstream analyses.

We performed a PCA analysis using the allele frequencies of all the SNPs to investigate the population structure using the package bigsnpr v.1.4.4^96^ in R, and found that the populations did not segregate into two separate genetic clusters (Supplementary Figure S4a-b). Further, we repeated this analysis for SNPs from regions under selection (see below) to test whether we observe the same population structure patterns. We observed that these separated into distinct resistant and susceptible groups, with the exception of a resistant population, BI, which clustered between the susceptible and other resistant populations (Supplementary Figure S4c). Thus for the purposes of this study, we dropped the BI population from further analysis.

### Selection analysis

We split the high confidence variant dataset obtained into ‘resistant’ and ‘susceptible’ variant dataset using vcf-subset of VCFtools v0.1.15^94^. The ‘resistant’ and ‘susceptible’ variant datasets contained 30 and 39 individuals, respectively (Table S1). We then used these datasets to calculate diversity and selection statistics G_ST_^97^, pi, Tajima’s D^98^, Fu and Way’s H^99^ using a custom script from ^100^ in a 300SNP window, for both the dataset. Furthermore, to detect hard sweep we phased the variants using beagle version 5.1^101^ which was then used to calculate the haplotype homozygosity statistic (H12, a measure of haplotype homozygosity that detects both hard and soft sweeps) using the scripts provided^102^. For regions above 95 percentile G_ST_, we calculated a composite rank based statistic (Md-rank-*P*) which was computed as the Mahalanobis distance on the negative log10 transformation of raw statistics into rank P-values^103^. This Md-rank-P was calculated using pi, Tajima’s D, Fu and Way’s H, and H12. To identify potential regions of selection we chose bins with greater than 95 percentile Md-rank-*P*.

### Linkage analysis

We calculated linkage disequilibrium (r^2^) at three different levels. First, to estimate the background genome-wide long-distance (and interchromosomal) linkage disequilibrium (ILD), we calculated r^2^ values for 5842 SNPs separated by at least 100kb using VCFtools v0.1.15^94^ (--thin 100000 --interchrom-hap-r2). Second, we estimated the r^2^ for SNPs separated by at least 1kb in and between broad regions (0.75Mb upstream and downstream) around the five focused regions (with G_ST_ > 0.39) under selection using VCFtools v0.1.15^94^. Lastly, since one would expect higher linkage between regions with high differentiation, we also randomly chose five regions with high differentiation (showing no signs of selection) of similar lengths as the five focused regions above and compared its linkage values to those regions.

### RNA-Seq

To identify transcripts associated with glyphosate resistance and its potential cost, we sequenced transcriptomes of 17 individuals belonging to four different treatments; resistant control (Rc), susceptible control (Sc), resistant herbicide sprayed (Rh), and susceptible herbicide sprayed (Sh). Each treatment had multiple individuals (Rc-2, Sc-2, Rh-6, Sh-7; Supplementary TableS2). The seeds were grown in a controlled environment (growth chamber) to reduce variation due to environmental differences. 20 days after planting, we sprayed glyphosate (concentration of 1.52 kg ai/ha) on the Rh and Sh treatment plants and collected the second and fourth leaf for RNA extractions 8 hours post-spray. These were flash frozen using liquid nitrogen and stored at -80°C. We extracted RNA using Qiagen RNeasy Plant mini kit with the optional DNase digestion step. This was then sequenced using Illumina NovaSeq 6000 at 150bp paired-end sequencing. A total of 132,551,535,000 bp were obtained.

### Differential gene expression

We processed the raw reads obtained to remove adapters using cutadapt v1.18^104^ and then mapped them to the de-novo assembled genome (-- sjdbOverhang 149 --outSAMtype BAM SortedByCoordinate Unsorted) using STAR v2.7.5^90^. Next, using HTSeq v0.11.1^105^, we counted read counts for each gene. These read counts were then used to filter out lowly expressed transcripts using the Bioconductor package edgeR version 3.18.1^106^ such that transcripts were retained only if they had greater than 0.5 counts-per-million in at least two samples (Rc vs Sc) and four samples (Rh vs Sh). The libraries were then normalized in edgeR (using the trimmed mean of M-values method) followed by differential gene expression analysis using the classic pairwise comparison of edgeR version 3.18.1. We extracted the significance of differentially expressed transcripts (DETs) with FDR <= 0.05. This was done for two contrasts, Rh vs Sh (total sample size = 13; Rh = 6, Sh = 7) and Rc vs Sc (total sample size 4; Rc = 2, Sc = 2). The first contrast informs us of the genes that are regulated in response to the herbicide, and how this gene regulation differs between the resistant and the susceptible populations, whereas the latter informs us of the baseline expression difference due to glyphosate resistance between the two populations.

### Malathion Experiment

On May 15th, 2019, we planted a total of 180 replicate seeds from multiple resistant and susceptible populations (Supplementary TableS3) in Cone-Tainers (Stewe and Sons). These were allowed to grow for 30 days, after which we subjected them to one of the four treatment environments--malathion (7.81 ml/L according to manufacturer’s recommendations), glyphosate (3.4 kg ai/ha), glyphosate and malathion, and a control. Twenty-five days post treatment spray, we recorded death as a metric trait (dead/almost dead or green and healthy), and harvested the plants. These were dried for 3 days at 70C and weighed for an estimate of dry above ground-biomass.

Using this data, we assessed whether biomass was significantly altered by the different treatments. First, we normalized the above-ground biomass using the transformTukey function from Rcompanion v.2.0.0^107^. We then used a generalized linear model (lm function^108^ with as normalized biomass as the dependent variable and population Type (R/S) and treatment as the independent variables. We assessed the significance of the variables using the Anova function of the car package v.3.0.10^109^, and performed a pairwise comparison between groups using the lsmeans function from package lsmeans v2.30.0^110^, adjusted for multiple tests using tukey correction. Using the same general model, we also compared whether biomass was significantly different between treatments for each population type. To control for the differences in the plant size we standardized the biomass of the individuals by the average biomass of the respective maternal line in the control treatment, and then normalized it as above.

## Data Availability

The datasets generated during and/or analysed during the current study have been deposited in GeneBank database under the project XXX, which are publicly accessible at XXX.

The *I. purpurea* genome assembly and annotation are available in the CoGe platform - https://genomevolution.org/coge/GenomeInfo.pl?gid=58735.

## Code Availability

The custom codes used in this study are deposited in GitHub (https://github.com/gsonal802/IP_GS.git).

## Acknowledgements

We thank M. Rausher for supplying germplasm used in the *I. purpurea* genome assembly, and T. Newsum, S. Paranjape, and K. Johnson for growth room phenotyping, and M. Palmer and MBGNA for growth room support. We likewise thank J. Opp in the University of Michigan sequencing core for sequencing support, Amanda Cummings for high molecular weight DNA preparation, and the Georgia Genomics and Bioinformatics Core, which provided the PacBio Sequel sequencing service. We also thank E.B. Josephs, R.L. Rogers, and the Ross-Ibarra lab for providing feedback on the manuscript. Funding for this work was provided by the University of Michigan and USDA NIFA awards 24892 & 28497 to RSB, NSF DEB 1442199 and IOS 1444567 to J.L.M., and NSF IOS 1611853 to A.H.

## Footnotes

^1^S.G. and A.H. contributed equally to this work.

^2^Author contributions: R.S.B., and S.G. designed the project; R.S.B., M.L.V.E., and S.G. designed the experiments; A.H. and J.L.M. assembled and annotated the genome; A.S. performed the growth room experiment; M.L.V.E. performed the WGS experiment, S.G. analyzed whole genome resequencing, RNAseq data, and performed linkage analysis; and R.S.B., S.G., A.H., and J.L.M. wrote the paper.

The authors declare no competing interest.

