## Supplementary Data for "Inter-chromosomal linkage disequilibrium and linked fitness cost loci associated with selection for herbicide resistance"

### Supplemental Figures

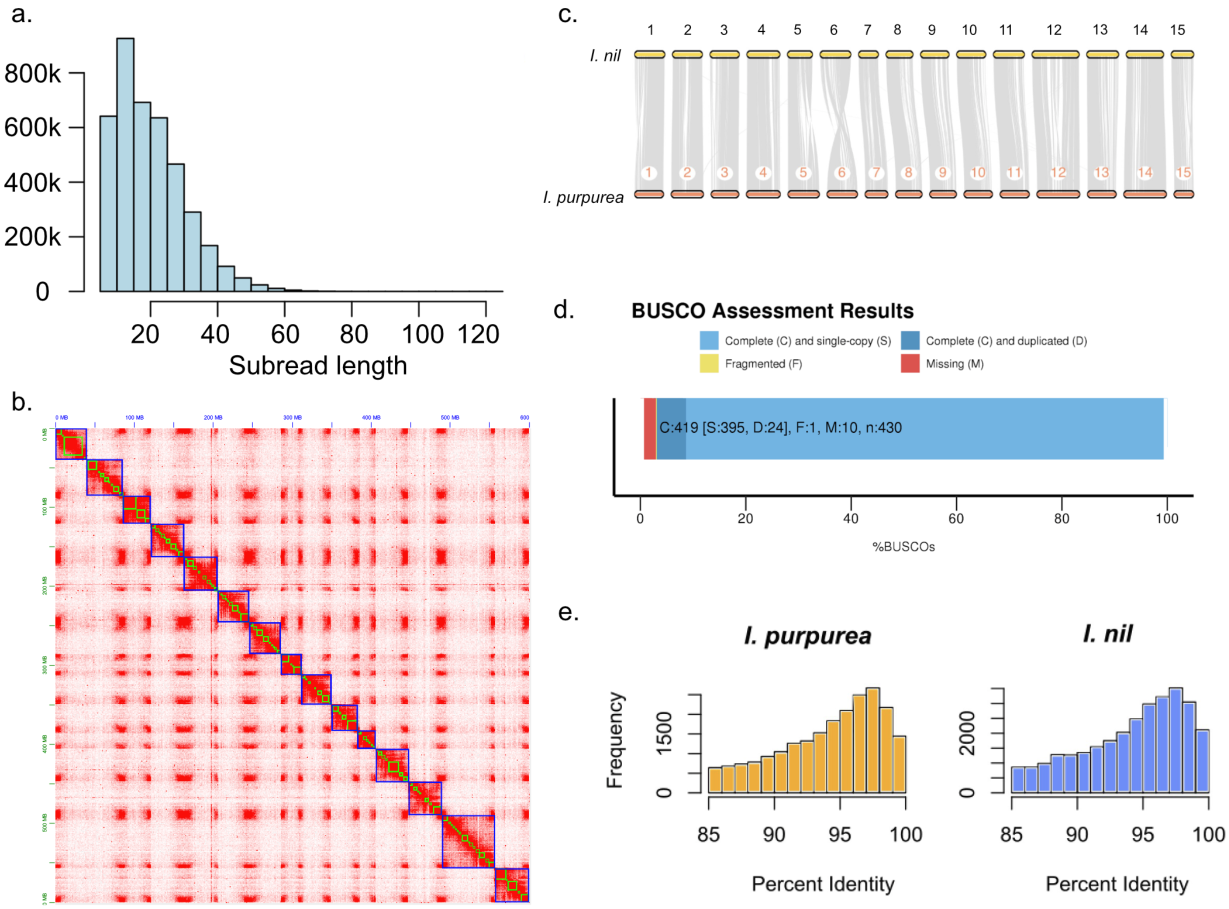


Supplementary Figure S1: Chromosome scaffolding and renaming. a) Raw PacBio Sequel filtered subread lengths. b) Phase Genomics Hi-C Proximo scaffolding results produces 15 chromosome pseudomolecules. c) Synteny of this *Ipomoea purpurea* assembly against *Ipomoea nil* was used to orient and name *I. purpurea* pseudomolecules. d) BUSCO results against Viridiplantae odb10 indicate high completeness of conserved gene sets in the raw assembly. e) Retrotransposon annotations using LTRharvest were used to compute pairwise LTR identities within LTRharvest for both *I. purpurea* and *I. nil*.

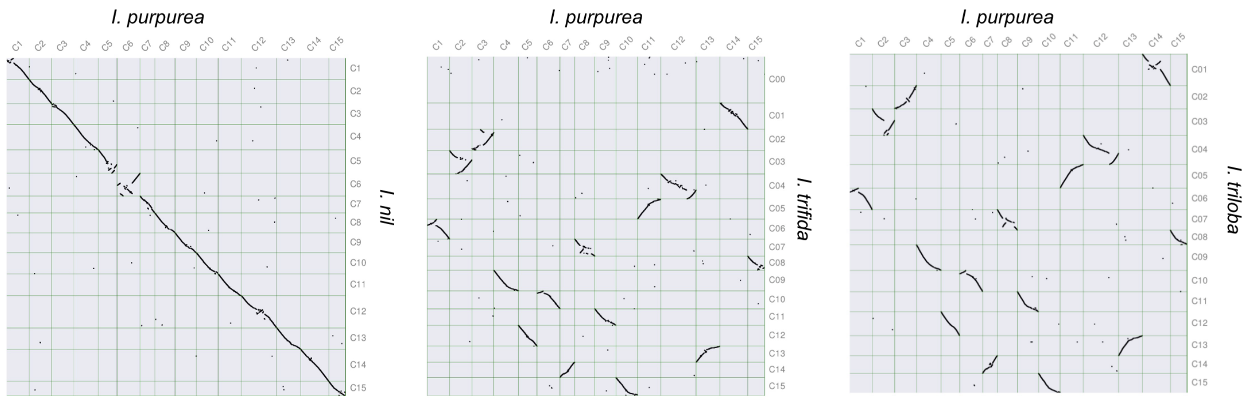


Supplementary Figure S2: Synteny of the *I. purpurea* genome against related Convolvulaceae species, including *I. nil*, *I. trifida*, and *I. triloba*.

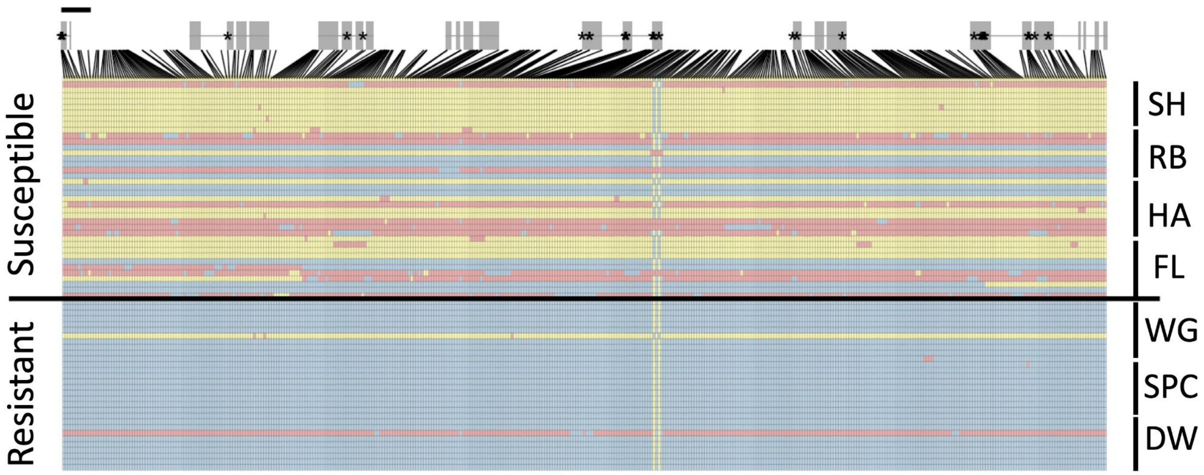


Supplementary Figure S3: Signs of selection across conserved haplotype of multiple glycosyltransferases for each individual on Chromosome10. Exons are shown in grey. Blue and yellow indicate homozygotes, red indicates heterozygotes; stars indicate non-synonymous substitutions. Black bar above gene models indicate 1kb.

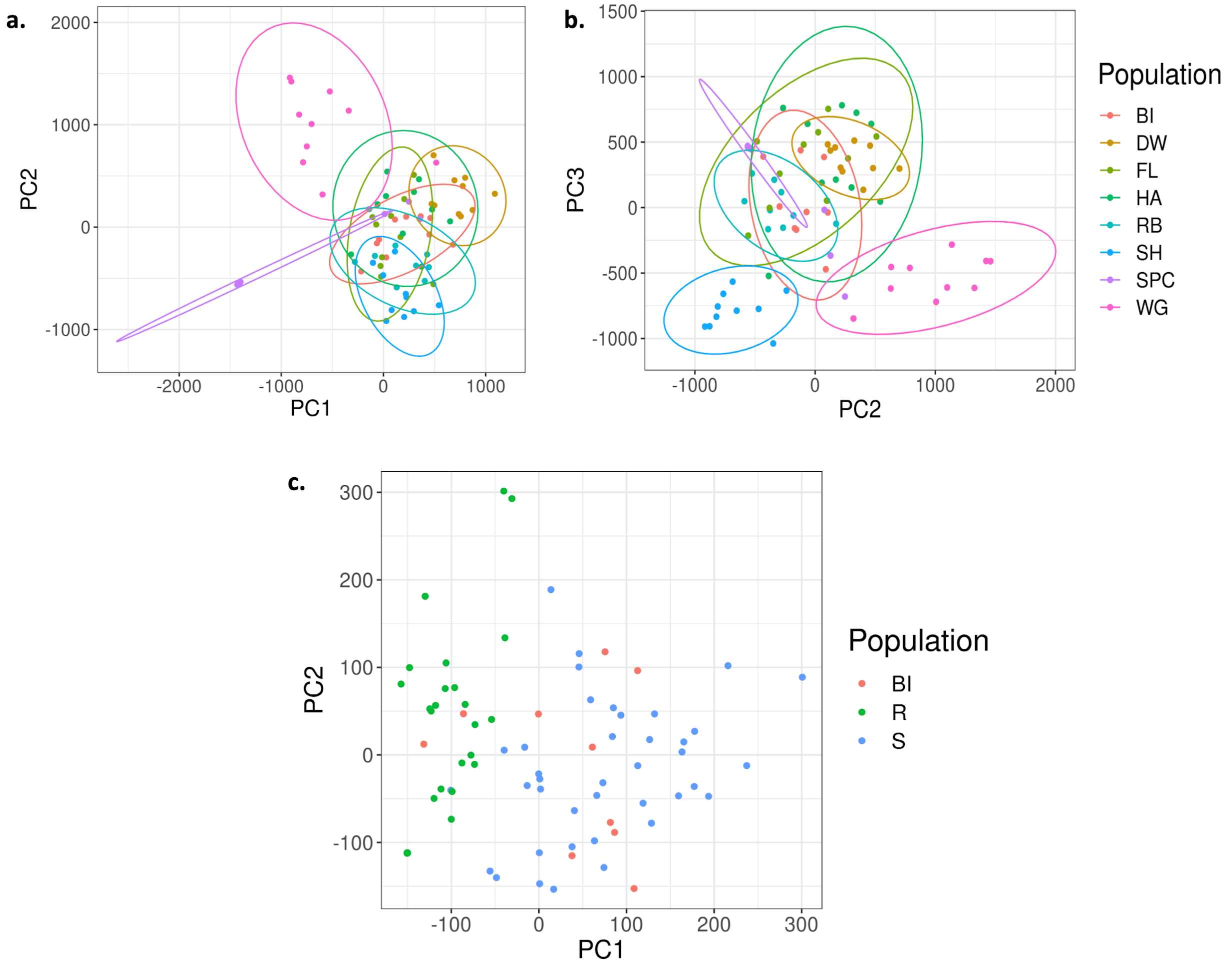


Supplementary Figure S4: Individuals from the sampled populations do not cluster into distinct resistant and susceptible groups when using all the SNPs (a and b), but there is some grouping when only considering the SNPs from the regions under selection (c).

]Table S1: Population information for each population used in the study.

| Pop Abbrev | Resistance Type | State | Proportion survival at 1.7 | Latitude | Longitude | No of individuals sampled |
|---|---|---|---|---|---|---|
| BI | R | TN | 1 | 35.775 | -85.903 | 10 |
| DW | R | NC | 1 | 34.983 | -78.039 | 10 |
| FL | S | SC | 0.20 | 34.145 | -79.865 | 10 |
| HA | S | NC | 0.15 | 35.424 | -77.917 | 10 |
| RB | S | TN | 0.18 | 35.316 | -87.353 | 9 |
| SH | S | VA | 0.1 | 38.373 | -78.662 | 10 |
| SPC | R | NC | 0.71 | 35.533 | -85.951 | 10 |
| WG | R | TN | 0.83 | 35.099 | -86.225 | 10 |

Pop Abbrev = abbreviation for each population as used in Kuester *et al.* 2015, Resistance type = classification of resistance in the population R >0.5 prop. survival S <0.5 prop. survival, State = state where seeds were collected, Proportion survival at 1.7 = proportion of individuals that survived a spray rate of 1.7 kg/ha of glyphosate based on Kuester et al 2015, Latitude and Longitude = location where seeds were collected.

Table S2: RNA-Seq sample information used in the study.

| Sample name | Pop Abbrev | Resistance Type | TRT |
|---|---|---|---|
| IP_438 | WG | R | Control |
| IP_447 | WG | R | Control |
| IP_235 | WG | R | Herbicide |
| IP_244 | DW | R | Herbicide |
| IP_247 | WG | R | Herbicide |
| IP_248 | WG | R | Herbicide |
| IP_252 | DW | R | Herbicide |
| IP_261 | WG | R | Herbicide |
| IP_459 | RB | S | Control |
| IP_477 | SH | S | Control |
| IP_177 | HA | S | Herbicide |
| IP_188 | RB | S | Herbicide |
| IP_189 | HA | S | Herbicide |
| IP_232 | HA | S | Herbicide |
| IP_257 | RB | S | Herbicide |
| IP_260 | RB | S | Herbicide |

| | | | |
|---|---|---|---|
| IP_497 | SH | S | Herbicide |

Pop Abbrev = abbreviation for each population as used in previous studies, Resistance type = classification of resistance in the population R >0.5 prop. survival S <0.5 prop. survival,TRT = Treatment type based on either herbicide sprayed (Herbicide) or not sprayed (Control).

Table S3: Sample information used for the malathion assay.

| Pop Abbrev | Resistance Type | TRT | Sample size |
|---|---|---|---|
| BI | R | Malathion | 6 |
| DW | R | Malathion | 13 |
| WG | R | Malathion | 22 |
| HA | S | Malathion | 4 |
| IN | S | Malathion | 3 |
| RB | S | Malathion | 11 |
| SH | S | Malathion | 3 |
| BI | R | Glyphosate | 3 |
| DW | R | Glyphosate | 5 |
| WG | R | Glyphosate | 8 |
| HA | S | Glyphosate | 3 |
| RB | S | Glyphosate | 4 |

| | | | |
|---|---|---|---|
| BI | R | Glyphosate + Malathion | 2 |
| DW | R | Glyphosate + Malathion | 9 |
| WG | R | Glyphosate + Malathion | 13 |
| HA | S | Glyphosate + Malathion | 4 |
| RB | S | Glyphosate + Malathion | 6 |
| BI | R | Control | 5 |
| DW | R | Control | 12 |
| WG | R | Control | 20 |
| HA | S | Control | 8 |
| IN | S | Control | 3 |
| RB | S | Control | 10 |
| SH | S | Control | 3 |

Pop Abbrev = abbreviation for each population as used in previous studies, Resistance type = classification of resistance in the population R >0.5 prop. survival S <0.5 prop. survival,TRT = Treatment type, Sample size = Total number of individuals per population per resistance type per treatment.

Table S4. Summary of the genome-wide regions under selection.
(Provided as a tabular data in excel file)

Table S5: Functional annotations of genes under selection
(Provided as a tabular data in excel file)

Table S6: List of differentially expressed genes between treated (herbicide sprayed) resistant vs susceptible individuals.
(Provided as a tabular data in excel file)

Table S7: List of differentially expressed genes between control (non-herbicide sprayed) resistant vs susceptible individuals.
(Provided as a tabular data in excel file)

Table S8: Pairwise contrast statistics for normalized above-ground biomass between the four treatment conditions. These were calculated using the lsmeans function in R, with P-values adjusted for multiple tests using tukey correction.

| Contrast | Contrast Estimate | t-ratio | P-value |
|---|---|---|---|
| Malathion vs Glyphosate | 0.455 | 3.913 | 0.0007 |
| Malathion vs Malathion-Glyphosate | 0.834 | 8.200 | <0.0001 |
| Malathion vs Control | -0.460 | -5.342 | <0.0001 |
| Glyphosate vs Malathion-Glyphosate | 0.379 | 2.946 | 0.0190 |
| Glyphosate vs Control | -0.915 | -7.834 | <0.0001 |
| Malathion-Glyphosate vs Control | -1.29 | -12.654 | <0.0001 |

Table S9: ILD summary statistic (99th percentile value and max $r^2$) for the five regions under selection that exhibited $G_{ST} > 0.39$. The summary is reported only for only the SNPs within regions under selection. The 99th percentile reports the top 1% of $r^2$ values whereas the max $r^2$ value is the highest $r^2$ value in the region.
(Provided as a tabular data in excel file)

Table S10: Individual ILD interactions, above the 99-percentile cutoff $r^2$ value, for SNPs within the region under selection.
(Provided as a tabular data in excel file)
